# Serotonergic innervation and cortical progenitor regulation in human brain assembloids

**DOI:** 10.64898/2026.04.13.718159

**Authors:** Raquel Pérez Fernández, Marco Torben Siekmann, Matteo Gasparotto, Lutz Wallhorn, Annasara Artioli, Martin Kubitschke, Catello Guida, Ammar Jabali, Anne Hoffrichter, Philipp Koch, Olivia Andrea Masseck, Julia Ladewig

## Abstract

Neuromodulatory signaling is classically associated with mature neural circuits, yet serotonergic projections reach the developing cortex prior to circuit formation, suggesting a role in early human cortical development. Here, we establish a human raphe–cortical assembloid platform by generating iPSC-derived hindbrain-patterned raphe organoids that produce serotonergic neurons and form projections into fused cortical organoids, exhibiting endogenous serotonin release within developing cortical tissue. Using this system, we find that serotonergic innervation is associated with a shift toward progenitor-enriched states, accompanied by increased proliferative activity, activation of developmental transcriptional programs, and predicted signaling interactions targeting cortical progenitors and neurons. Consistent with these findings, cortical regions receiving serotonergic projections exhibit increased mitotic activity, and pharmacological modulation demonstrates selective proliferative responses in basal progenitor populations, consistent with observations in human fetal tissue. Together, this system provides a framework to investigate how early neuromodulatory input shapes human cortical development and developmental vulnerability.

## Introduction

Neuromodulatory systems are traditionally studied in the context of mature brain circuits, where they regulate neuronal activity and plasticity^1^. However, these systems emerge much earlier during brain development^2^, positioning them to influence fundamental cellular processes during neural tissue formation. This early presence is particularly relevant given increasing evidence linking dysregulation of neuromodulatory signaling to neuropsychiatric disorders such as autism spectrum disorders, anxiety disorders, and major depression, many of which originate during early developmental stages^3–5^.

Among neuromodulatory systems, the serotonergic system is especially well positioned to act during these early phases, contributing to the regulation of neural plasticity and network formation^6,7^. Serotonin is initially provided by maternal and placental compartments during early development^8^. Fetal endogenous serotonergic neurons arise in the raphe nuclei around gestational week 5^9^ and extend projections to the developing forebrain by gestational week 8^10^, coinciding with key periods of cortical progenitor expansion and highlighting sertonińs potential to influence fetal brain development at these stages^11^. Experimental studies in rodent models demonstrate that serotonergic signaling can modulate cortical progenitor dynamics^12–14^. These findings support a role for serotonin as an early modulatory input during cortical development.

Human corticogenesis represents a complex developmental process, characterized by extensive progenitor expansion and lineage diversification^15^. Neural progenitors in the ventricular and subventricular zones (VZ and SVZ) undergo tightly regulated transitions that determine cortical growth and neuronal output^16,17^. In humans, basal progenitors—particularly basal radial glia—are markedly expanded and are thought to underlie the evolutionary enlargement of the cortex compared to lissencephalic species such as rodents^18–20^. Importantly, components of serotonergic signaling also exhibit species-specific patterns, with receptors such as HTR2A expressed in the developing neocortex of humans and ferrets but largely absent in the mouse^21,22^. Consistent with this, serotonergic signaling has been shown to selectively influence progenitor behavior in the developing human cortex, including effects on radial glia function and basal progenitor proliferation^21,22^. These features highlight the importance of studying early developmental regulation in human systems.

However, investigating how neuromodulatory inputs influence cortical development in this context remains challenging. Existing approaches largely rely on exogenous manipulation of isolated tissue systems^21,22^, which fail to capture endogenous long-range input to developing cortical tissue. Moreover, access to primary human fetal tissue is limited, and experimental manipulation remains constrained. Human stem cell–derived organoids and assembloids have emerged as powerful systems to model early brain development and to study complex processes such as corticogenesis^23,24^ and interregional interactions, including interneuron migration and circuit-level connectivity^25–27^. Notably, systems that combine physiologically relevant neuromodulatory projections with the ability to resolve responses at the level of cortical progenitor populations remain limited.

Here, we developed an iPSC-derived raphe–cortical (RO–CO) assembloid system to model early serotonergic input during human cortical development. Hindbrain-patterned raphe organoids give rise to serotonergic neurons that release serotonin and project into fused cortical organoids. Integration of serotonergic input in this system is associated with shifts in cortical progenitor states, including increased proliferative activity and activation of developmental transcriptional programs. In line with these changes, innervated cortical regions exhibit elevated mitotic activity, while pharmacological perturbation indicates selective responsiveness of basal progenitor populations.

Together, these findings establish a human assembloid platform for studying early neuromodulatory influences on cortical development and demonstrate that serotonergic input can be functionally integrated to modulate cortical progenitor dynamics in vitro. This approach provides a framework to investigate how neuromodulatory signaling shapes early developmental trajectories and contributes to developmental vulnerability.

## Results

### Generation and characterization of raphe organoids from iPSCs modeling serotonergic identity

To model early serotonergic input in a human system, we generated hindbrain-patterned organoids predominantly resembling the serotonergic raphe nuclei (ROs; **Figure 1A**).

**Figure 1.**
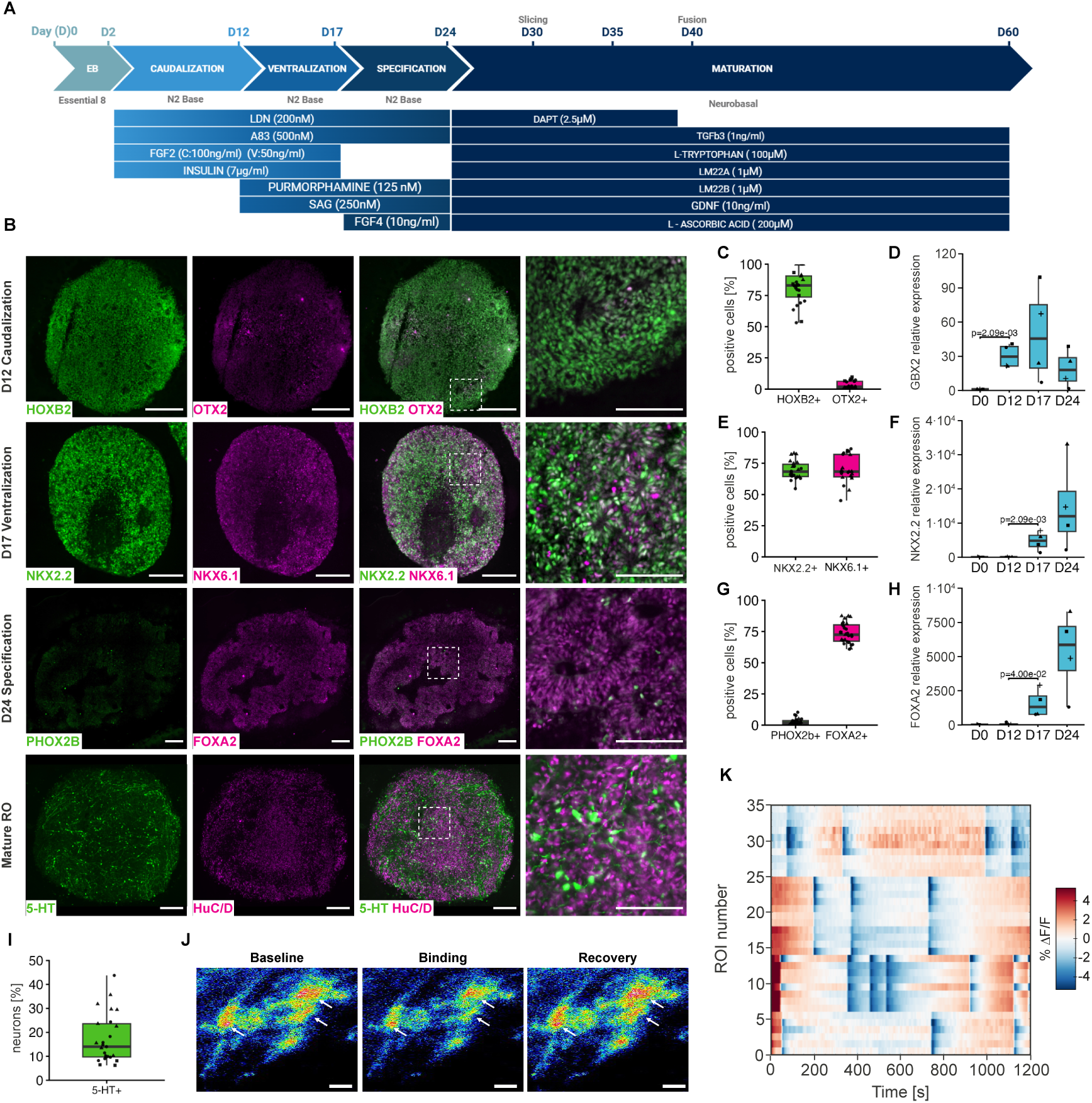
Generation and characterization of raphe organoids. **A** Schematic overview of the differentiation protocol for hindbrain-patterned raphe organoids (ROs), indicating stage-specific signaling factors driving caudalization, ventralization, progenitor specification, and neuronal maturation. **B** Representative immunofluorescence images of ROs at sequential stages of differentiation showing marker expression during caudalization (day 12; HOXB2, green; OTX2, magenta), ventralization (day 17; NKX2.2, green; NKX6.1, magenta), serotonergic lineage specification (day 24; PHOX2B, green; FOXA2, magenta), and neuronal maturation (day 60; 5-HT, green; HuC/D, magenta). White boxes indicate higher-magnification regions. **C**–**I** Quantification and RT–qPCR analysis of stage-specific marker expression across differentiation stages. Quantification of marker expression (C, E, G, I). Individual data points represent measurements from organoid sections (fields of view), expressed as the percentage of marker-positive cells relative to total cells (DAPI+; C, E, G) or neurons (HuC/D+; I). Day 12: HOXB2+, OTX2+ (n = 24, 7 batches, 3 lines; C); day 17: NKX2.2+, NKX6.1+ (n = 22, 6 batches, 2 lines; E); day 24: PHOX2B+, FOXA2+ (n = 27, 7 batches, 3 lines; G); day 45: 5-HT+ (n = 25, 6 batches, 3 lines; I). RT–qPCR analysis during differentiation (D, F, H; day 0, 12, 17, 24). Expression of the hindbrain marker GBX2 (D), the ventral p3 domain marker NKX2.2 (F), and the floor plate marker FOXA2 (H) increased across stages (n = 4 batches). Boxplots (C-I) show the median, interquartile range (IQR, 25th–75th percentiles), and whiskers extending to 1.5× IQR. Statistical analysis (D, F, H) was performed using two-sided Student’s t-tests comparing each stage to the preceding time point, with Benjamini–Hochberg correction. **J-K** Functional assessment of serotonergic activity using the genetically encoded serotonin sensor sDarken. Representative pseudocolor images of sDarken-expressing cells showing baseline fluorescence, decreased fluorescence during endogenous serotonin signaling, and subsequent recovery of fluorescence signal. White arrows indicate regions exhibiting dynamic changes in fluorescence intensity (J). Heatmap of fluorescence changes (% ΔF/F) across 35 regions of interest (ROIs) over time from 4 independent batches derived from 1 line (K). Scale bars: B, overview 200 µm; high magnification 100 µm; J, 10 µm.

Serotonergic neurons arise from progenitors in the ventral p3 domain of the hindbrain, specified by coordinated rostrocaudal and dorsoventral patterning cues^7,28^. To induce and maintain neuroectodermal fate, we applied dual SMAD inhibition^29^. Caudalization of iPSC aggregates with FGF2- and insulin^30^ established a posterior hindbrain identity by day 12, as indicated by widespread HOXB2 expression and minimal OTX2 signal (**Figure 1B, C**). Consistently, expression of the hindbrain marker *GBX2* increased during early differentiation and remained elevated throughout subsequent patterning stages (**Figure 1D**). Subsequent activation of Sonic Hedgehog (SHH) signaling using puromorphamine and SAG^31^ directed progenitors toward a ventral p3 domain identity, marked by NKX2.2 and NKX6.1 co-expression (**Figure 1B, E, F**). Within this domain, FGF4-mediated patterning biased ventral progenitors toward a serotonergic lineage, characterized by strong FOXA2 expression and minimal PHOX2B, indicating selective enrichment of serotonergic progenitors over alternative fates. (**Figure 1B, G, H**). Upon neuronal maturation (> day 45), ROs contained serotonergic neurons, identified by co-expression of HuC/D and 5-HT (**Figure 1 B, I**). Expression of the serotonin-synthesizing enzyme TPH2 together with the vesicular monoamine transporter VMAT2 further confirmed serotonergic specification (**Supp. Fig. S1A, B**). RO identity and serotonergic differentiation were consistently observed across independent differentiations and iPSC lines (**Figure 1C, E, G, I; Supp. Fig. S1C**).

To assess serotonin dynamics, we employed the genetically encoded serotonin sensor sDarken^32^. RO slices maintained in air–liquid interface culture^33^ and expressing sDarken displayed spontaneous transient decreases in fluorescence across the tissue, consistent with serotonin release and reuptake dynamics, including recovery characteristics indicative of serotonin clearance (**Figure 1J, K; Supp. Fig. S1D, E**). Similar fluorescence responses were observed in neurons derived from dissociated ROs expressing sDarken, demonstrating that individual serotonergic neurons are capable of serotonin release independent of a network context (**Supp. Fig. S1F, G**).

These results establish ROs as a reproducible model for serotonergic signaling, demonstrating serotonin release in vitro and providing a platform to study early neuromodulatory signaling.

### Raphe–cortical assembloids establish serotonergic innervation and display a defined cellular composition

To model serotonergic input to the developing cortex, we generated RO-CO assembloids by fusing ROs with cortical organoids (COs)^34–36^ between days 35 and 45 of differentiation. Brightfield imaging revealed progressive structural integration of the two compartments over three weeks (**Figure 2A, left**). Immunofluorescence analysis demonstrated serotonergic projections extending from the raphe compartment into cortical tissue, indicating establishment of long-range serotonergic innervation (**Figure 2A, right**). The cortical identity of the target compartment was confirmed by expression of dorsal telencephalic progenitor and neuronal markers, including FOXG1, TBR1, and CTIP2 (**Figure 2B**). High-resolution imaging further detected punctate localization of synapsin and homer at serotonergic axon termini in proximity to the serotonin receptor HTR2A within cortical target regions, supporting potential synaptic contact formation between raphe-derived axons and cortical cells (**Supp. Fig. S2A**).

**Figure 2.**
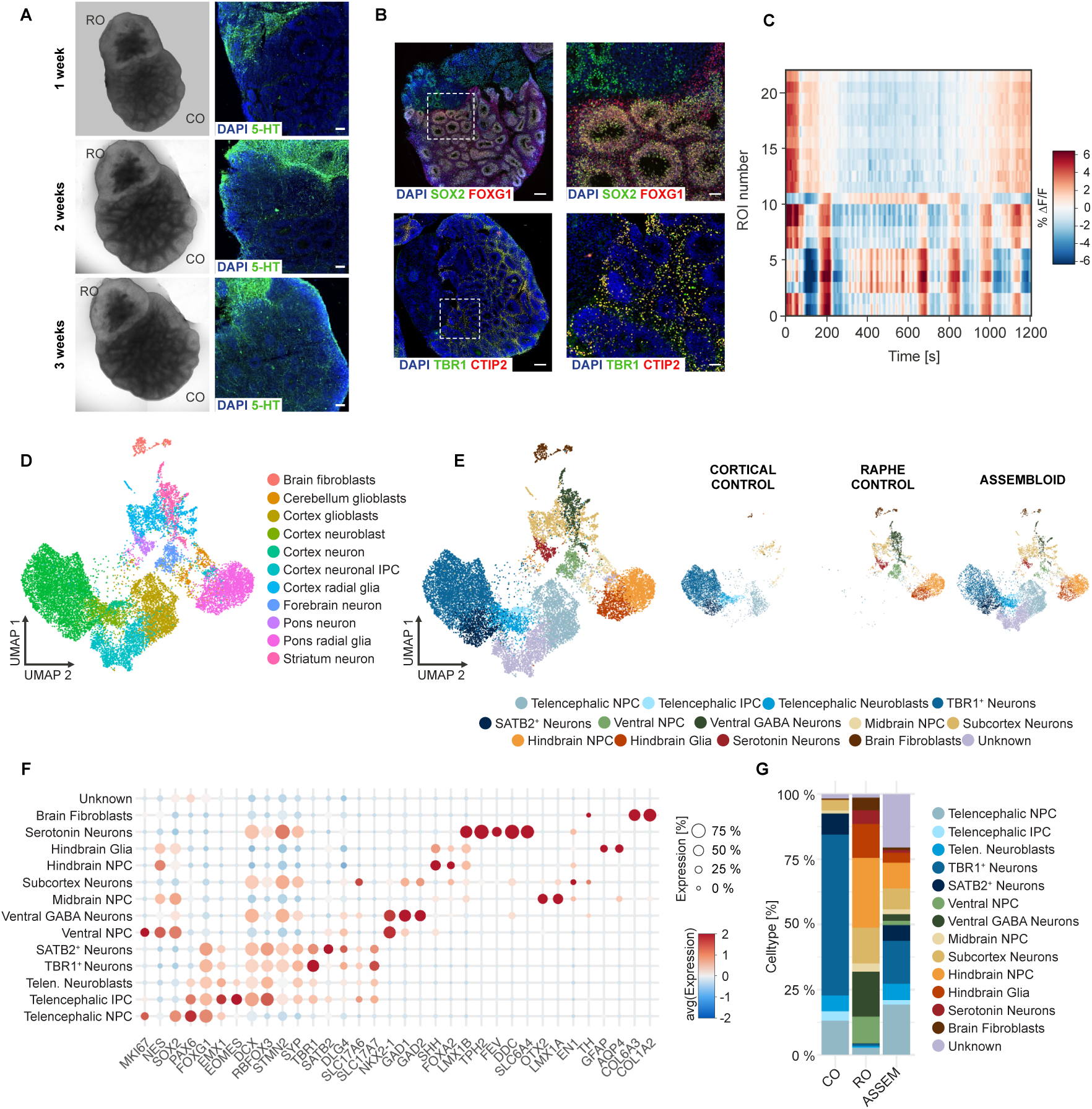
Establishment and characterization of raphe–cortical assembloids. **A** Representative bright-field images of raphe–cortical (RO-CO) assembloids at 1, 2, and 3 weeks after fusion, suggesting progressive structural integration of both compartments (left). Corresponding immunofluorescence images at the same time points show serotonergic projections (5-HT, green) extending from the raphe compartment into the cortical compartment (right; Counterstained with DAPI, blue). **B** Immunofluorescence staining of RO-CO assembloids (day 54) for cortical progenitor markers (FOXG1, red; SOX2, green; top) and cortical layer markers (TBR1, green; CTIP2, red; bottom) consistent with preservation of cortical identity. White boxes indicate higher-magnification regions. Counterstained with DAPI (blue). **C** Heatmap of fluorescence changes (% ΔF/F) in cortical regions of RO-CO assembloids across 22 regions of interest (ROIs) over time from 2 independent batches, showing dynamic fluorescence fluctuations consistent with endogenous serotonergic signaling. **D** UMAP visualization of snRNA-seq data from CO, RO, and RO-CO assembloids using CellTypist reference-based label transfer, showing assignment to telencephalic and hindbrain identities. **E** UMAP visualization identifying 14 transcriptionally distinct cell populations (left). UMAP plots split by sample (CO, RO, and RO-CO assembloids) showing distribution of cortical-and raphe-derived cell populations (right). **F** Dot plot of canonical marker gene expression across cell populations. Dot size indicates fraction of expressing cells and color represents scaled average expression levels. **G** Cell-type distribution across samples, showing relative proportions (%) of telencephalic and hindbrain-derived populations in CO, RO, and RO-CO assembloids. All snRNA-seq analyses shown in D–G are based on day 61 samples (9 organoids/assembloids; total n = 17,805 nuclei after quality filtering). Scale bars: A, 100 µm; B, overview 200 µm; high magnification 50 µm.

To assess serotonin dynamics within COs in the assembloids, we used the sDarken system^32^. Sensor responsiveness in a cortical context was first validated in iPSC-derived cortical neurons by exogenous serotonin application (**Supp. Fig. S2B**). In RO-CO assembloids, sDarken-expressing cortical regions exhibited recurrent fluorescence decreases in the absence of exogenous serotonin, demonstrating that cortical cells are capable of serotonin uptake from RO-derived projections. Characteristic recovery dynamics consistent with serotonin clearance and reuptake, further confirmed serotonergic input to cortical tissue (**Figure 2C, Supp. Fig. S2C**).

To define the cellular composition and transcriptional landscape of RO-CO assembloids, we performed snRNA-seq following 21 days of air–liquid interface culture and compared them to age-matched COs and ROs at day 61. After quality filtering, a total of 17,805 nuclei were retained for downstream analysis. An initial reference-based mapping using CellTypist^37^, based on a human developmental brain atlas^38^, identified both telencephalic and hindbrain populations (**Figure 2D**). Unsupervised clustering and UMAP visualization resolved 14 transcriptionally distinct populations (**Figure 2E**). These clusters were subsequently annotated based on established marker gene expression, revealing major dorsal telencephalic populations, including telencephalic neural progenitor cells (telencephalic NPCs: *SOX2*⁺, *PAX6*⁺, *FOXG1*⁺), intermediate progenitors (*EOMES*⁺), neuroblasts (*DCX*⁺), and excitatory cortical neurons corresponding to deep-layer (*TBR1*⁺) and upper-layer (*SATB2*⁺) identities. In contrast, ROs displayed hindbrain-derived populations dominated by hindbrain progenitors and glial-like cells (*SOX2*⁺, *SHH*⁺, *FOXA2*⁺, *LMX1B*⁺, *AQP4*⁺), as well as a distinct serotonergic neuronal cluster expressing canonical markers including *TPH2*⁺, *FEV*⁺, *DDC*⁺, and *SLC6A4*⁺. Additional clusters included *NKX2.1*-expressing ventral progenitors and their derived inhibitory neurons. Smaller fractions of subcortical neurons, midbrain-like progenitors, and fibroblast-like populations were also detected. A transcriptionally sparse cluster with low transcript counts and weak lineage marker expression remained unassigned (Unknown) (**Figure 2F**). This marker profile reveals that RO-CO assembloids contain contributions from both CO and RO identities, including dorsal telencephalic progenitors together with raphe-derived serotonergic neurons and hindbrain progenitors.

Together, these observations indicate that raphe-derived projections provide serotonergic input to cortical tissue within a cellularly integrated hindbrain–cortical assembloid system.

### Serotonergic input reshapes telencephalic NPC states and transcriptional programs

To determine how serotonergic input is associated with changes in cortical cell states, we analyzed telencephalic populations in RO-CO assembloids compared to CO controls. Cell-type proportion analysis revealed a marked shift from neuronal toward progenitor-enriched states within these populations (**Figure 2G, Supp. Table TS1, 2**). In COs, excitatory neurons (TBR1⁺ and SATB2⁺) accounted for ∼75% of telencephalic cells, whereas telencephalic NPCs represented ∼14%. In contrast, RO-CO assembloids showed a redistribution toward progenitor-enriched states, with telencephalic NPCs accounting for ∼40%. Intermediate progenitors remained relatively stable, while neuroblasts showed a relative increase. Within the neuronal compartment, this shift was primarily driven by a reduction in TBR1⁺ neurons, whereas SATB2⁺ neurons exhibited a modest increase.

To assess whether this shift is associated with changes in progenitor dynamics, we performed cell-cycle inference using Tricycle^39^. This analysis revealed an increased representation of S-phase cells among telencephalic NPCs in RO-CO assembloids compared to CO controls (**Figure 3A**), indicating increased proliferative activity.

**Figure 3.**
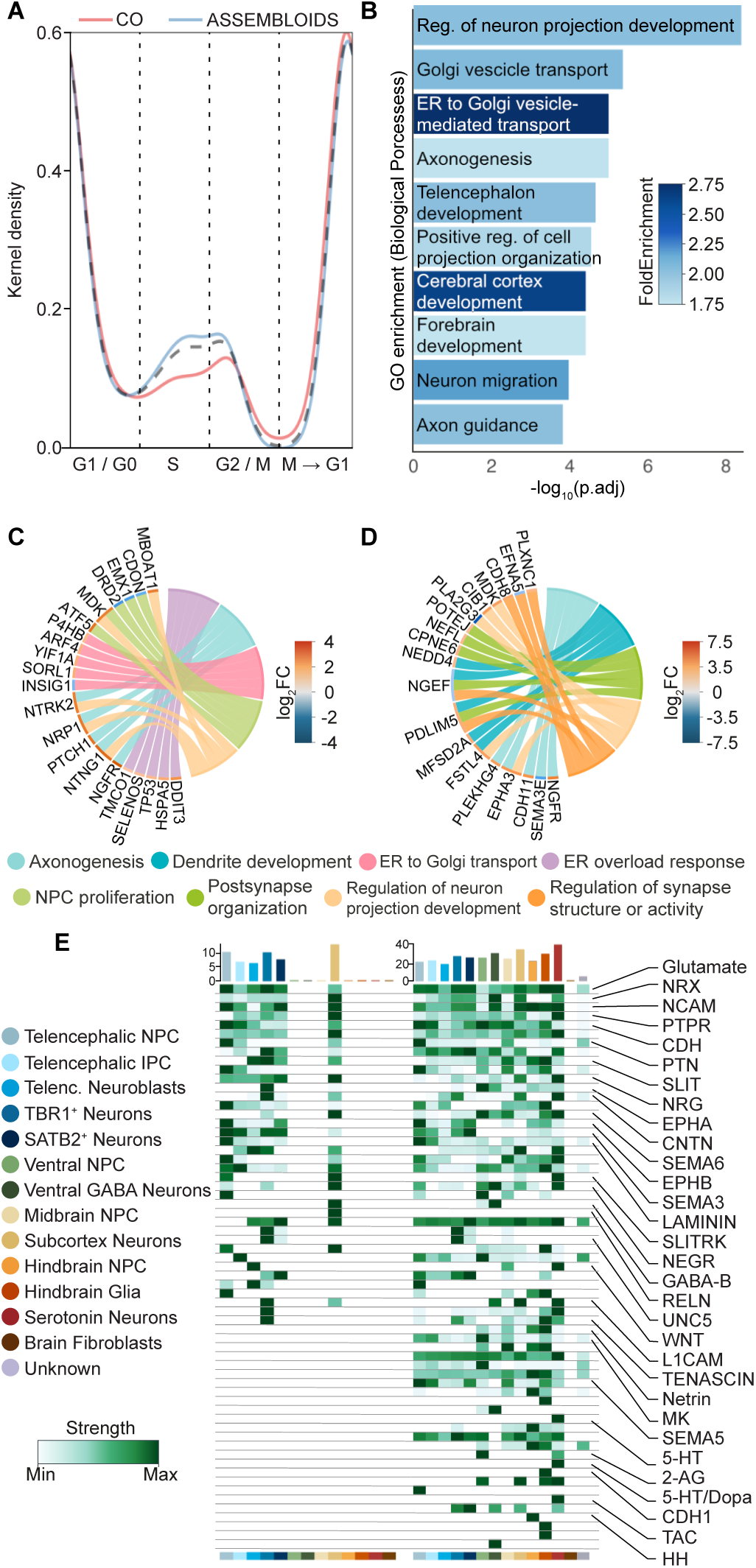
Serotonergic integration reshapes transcriptional programs and intercellular signaling in cortical populations. **A** Kernel density plot of inferred cell-cycle states (Tricycle) showing the distribution of telencephalic neural progenitors (NPC) across G1/G0, S, and G2/M phases in cortical organoids (COs) and raphe–cortical (RO-CO) assembloids. Telencephalic NPC in RO-CO assembloids show increased density in S-phase compared to COs. **B** Gene Ontology Biological Process (GO:BP) enrichment analysis of differentially expressed genes in telencephalic NPC upon fusion. Top 15 biological processes are shown, ranked by adjusted p-value. **C**–**D** Chord plots linking differentially expressed genes to enriched GO categories in telencephalic NPC (C) and TBR1⁺ neurons (D). **E** Heatmap of outgoing signaling patterns inferred from CellChat analysis comparing COs and RO-CO assembloids. Rows represent signaling pathways and columns represent sender cell populations. Color intensity indicates the strength of inferred signaling. The top bar plot shows the contribution of each cell population to the overall outgoing signaling network, with serotonergic neurons exhibiting the highest outgoing signaling activity. A subset of pathways is shown for visualization purposes (full dataset provided in Supplementary Tables 7-9). All snRNA-seq analyses shown in A–E are based on day 61 samples (9 organoids/assembloids; total n = 17,805 nuclei after quality filtering).

To define transcriptional programs associated with serotonergic input after fusion, we performed gene ontology (GO) enrichment analysis of differentially expressed genes across telencephalic populations. In telencephalic NPCs, enriched biological processes were dominated by developmental and morphogenetic programs related to neuronal differentiation and projection formation, together with intracellular trafficking and secretory pathways (**Figure 3B, Supp. Table TS3, 4**). In contrast, neuronal populations showed enrichment for pathways associated with morphogenesis and synaptic organization (**Supp. Fig. S3A, Supp. Table TS5, 6**). Analysis of associated gene sets highlighted genes linked to neuronal development and projection-related processes, including NRP1, NTNG1, NTRK2, and MDK, as well as genes involved in intracellular trafficking and secretory pathways such as SORL1, ARF4, and YIF1A (**Figure 3C, Supp. Table TS3, 4**). In neuronal populations, genes associated with morphogenesis and synaptic organization included *SEMA3E*, *EPHA3*, and *PLXNC1* (**Figure 3D, Supp. Table TS5, 6**).

To examine changes in intercellular communication upon assembly, we applied CellChat^40^ to infer signaling networks across cell populations. Compared to COs alone, RO-CO assembloids displayed a markedly expanded signaling landscape across cortical populations, while serotonergic neurons emerged as the cell population with highest outgoing signaling activity, identifying them as a major predicted signaling hub within fused systems. Outgoing signaling from serotonergic neurons was enriched for neuromodulatory pathways, including 5-HT, serotonin/dopamine, TAC, and 2-AG signaling, alongside broader developmental communication pathways such as NRG, CNTN, and L1CAM (**Figure 3E, Supp. Tables TS7-9**). Pattern analysis further organized serotonergic neuron–derived outgoing signaling into coordinated modules, with pattern 2 capturing a subset of these neuromodulatory and metabolic pathways (including 5-HT, TAC and cholesterol-related signaling; **Supp. Fig. S3B**). Predicted interactions were directed toward multiple cortical populations, including telencephalic NPCs, intermediate progenitors, and early excitatory neurons (TBR1⁺ and SATB2⁺; **Supp. Fig. S3C**), consistent with broad serotonergic communication across the developing cortical compartment. Analysis of incoming signaling further indicated a markedly expanded signaling landscape targeting cortical populations in RO-CO assembloids compared to controls **(Supp. Fig. S3D, Supp Table TS7, TS10,11)**. In telencephalic NPCs, this included broader network-level changes as well as a subset of newly detected pathways that were compatible with the overall signaling profile of RO-derived cell populations.

Together, these findings indicate that serotonergic input promotes a shift toward progenitor-enriched states and increased proliferative activity, accompanied by coordinated changes in transcriptional and signaling programs in telencephalic NPCs.

### Serotonin promotes cortical progenitor proliferation and selectively regulates basal progenitors via HTR2A signaling

SnRNA-seq of RO-CO assembloids revealed a shift toward progenitor-enriched states within the cortical compartment, accompanied by increased S-phase representation in telencephalic NPCs. In parallel, transcriptional analyses identified enrichment of developmental and morphogenesis-related programs, together with intracellular trafficking pathways.

To determine whether serotonergic projections directly influence progenitor dynamics at the tissue level, we quantified mitotic activity within cortical VZ/SVZ structures receiving these afferents. Immunofluorescence analysis revealed a significant increase in mitotic progenitors specifically in innervated regions (**Figure 4A, B**).

**Figure 4.**
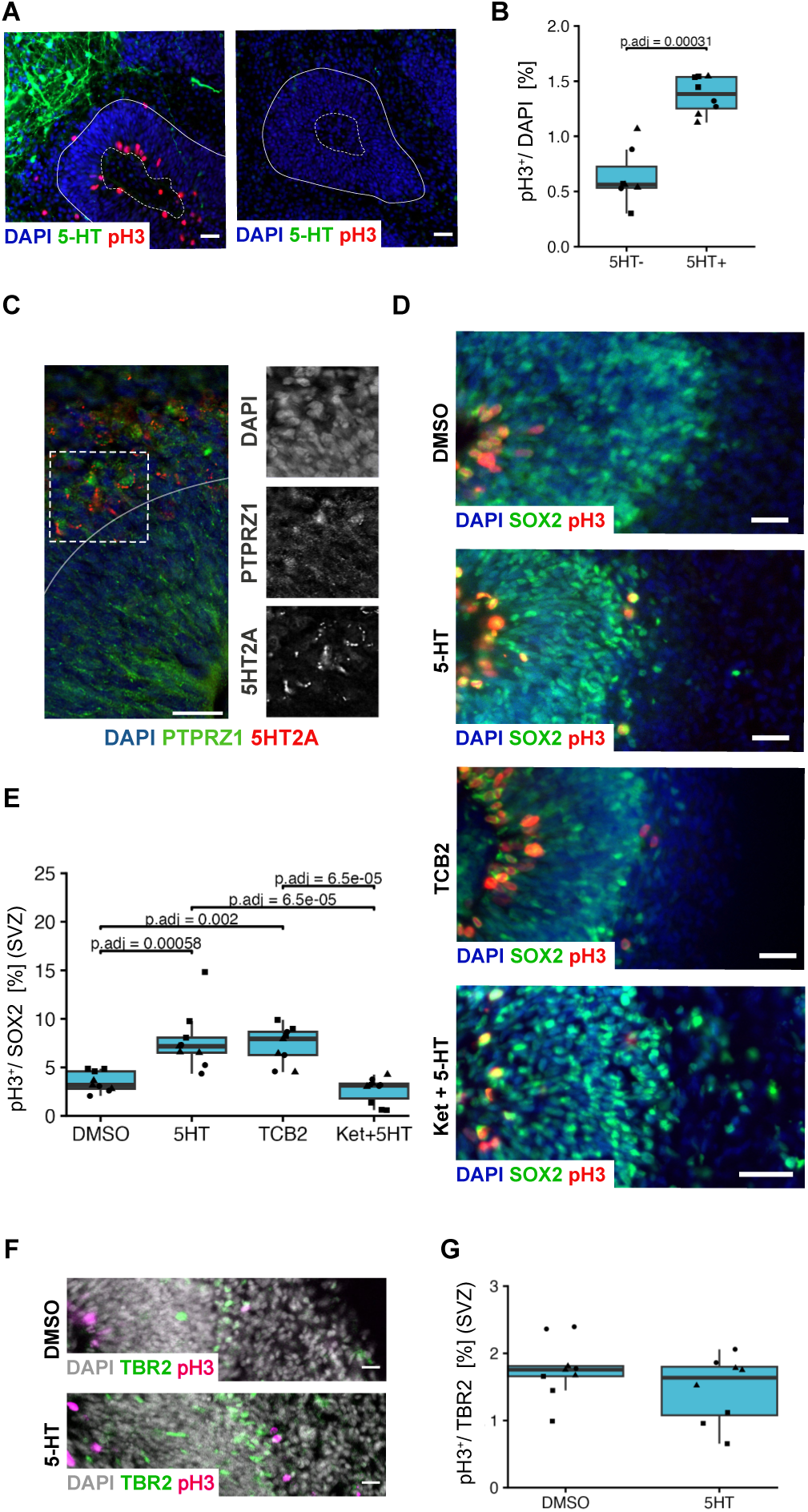
Serotonergic signaling regulates cortical progenitor proliferation via receptor-specific mechanisms. **A** Representative immunofluorescence images of raphe–cortical (RO–CO) assembloids showing cortical ventricular zones (VZ; DAPI, blue), with apical surface indicated by a white dotted line and the basal surface indicated by a white line, in regions with (left) or without (right) serotonergic projections (5-HT+, green) and mitotic cells (pH3+, red). **B** Quantification of pH3+ mitotic cells in cortical VZ/SVZ of RO–CO assembloids, comparing the regions with and without serotonergic innervation. A total of 73 VZ/SVZ structures were analyzed and aggregated into n = 15 independent regions for statistical analysis (3 batches, 1 line). **C** Representative immunofluorescence images of day 60 cortical organoids (COs) showing HTR2A receptor distribution (red, co-stained with PTPRZ1, green), counterstained with DAPI (blue). White lines indicate basal VZ surface. Inset indicate higher-magnification shown on the right with individual channels displayed in greyscale. **D** Representative immunofluorescence images of pH3 (red) and SOX2 (green) in day 60 COs following five days of treatment (DMSO, 5-HT, TCB2, Ketanserin + 5-HT), counterstained with DAPI (blue). **E** Quantification of basal progenitor proliferation, measured as the proportion of pH3^+^ SOX2^+^ cells in the SVZ. A total of 331 SVZ regions were analyzed (total n = 37 organoids; DMSO: n = 9; 5-HT: n = 9; TCB2: n = 9; Ket + 5-HT: n = 10; 3 batches). **F** Representative immunofluorescence images of pH3 (magenta) and TBR2 (green) in day 60 COs following five days of treatment (DMSO; 5-HT). **G** Quantification of intermediate progenitor proliferation, measured as the proportion of pH3⁺ TBR2⁺ cells (total n = 17 organoids; DMSO: n = 9; 5-HT: n = 8 organoids, 3 batches). B, F, H Boxplots show median, interquartile range (IQR; 25th–75th percentiles), and whiskers extending to 1.5× IQR. Individual points represent independent regions or organoids and are shaped by batch. Statistical analysis was performed using two-sided Wilcoxon rank-sum tests with Benjamini–Hochberg correction. Adjusted p-values are indicated where significant. Scale bars: A, 20 µm; C,D 25 µm; F, 15 µm.

Given prior evidence implicating HTR2A signaling in the regulation of basal progenitor proliferation in human fetal cortex^21^, we examined its distribution across cortical progenitor zones in COs. Immunostaining revealed enrichment of HTR2A in the SVZ relative to the VZ (**Figure 4C**). To functionally test receptor-specific effects, we performed controlled pharmacological perturbations in COs at day 60, corresponding to the developmental stage analyzed in the snRNA-seq dataset (day 61). Serotonin increased mitotic activity among SOX2⁺ basal progenitors. This effect was recapitulated by activation of HTR2A using the selective agonist TCB2 and attenuated by its inhibition with ketanserin, supporting HTR2A-dependent regulation of basal progenitor proliferation (**Figure 4D, E**). In contrast, TBR2⁺ intermediate progenitors did not exhibit significant changes in proliferation following serotonergic stimulation (**Figure 4F, G**), consistent with their stable representation in the snRNA-seq data (**Figure 2G**).

Together, these findings indicate that serotonergic input increases cortical progenitor proliferation and selectively engages basal progenitor populations via HTR2A-dependent mechanisms.

## Discussion

Neuromodulatory signaling has been primarily studied in the context of mature neural circuits^1^. Here, we establish a human iPSC–derived raphe–cortical assembloid system that enables the investigation of early neuromodulatory interactions in a controlled human context. Within this system, serotonergic innervation is associated with a shift toward progenitor-enriched states and increased proliferative activity, indicating that neuromodulatory input intersects with early developmental processes, including progenitor maintenance and lineage progression, prior to functional network formation. At the transcriptional level, these changes involve gene programs linked to neuronal development and projection-related processes, which are tightly regulated during corticogenesis^41^. In addition, the enrichment of intracellular trafficking and secretory pathways may reflect an increased responsiveness of cortical cells to incoming signals^42^. It will be important to determine how these transcriptional alterations are established and whether they reflect longer-term changes in cellular state. These effects may arise through downstream signaling, transcriptional regulation, or epigenetic mechanisms, and will help define how early neuromodulatory input influences developmental progression^43^.

A key feature of the assembloid system is that it capture early stages of human brain development, providing experimental access to developmental windows that are otherwise difficult to study in primary human tissue^44,45^. In addition, this system enables the investigation of neuromodulatory input through endogenous projections in a spatially and temporally controlled manner within a defined interregional context, which cannot be recapitulated by exogenous ligand application or isolated organoid systems. This is particularly relevant in the context of neuromodulatory signaling, where receptor expression and progenitor responsiveness appear to change dynamically over developmental time. For example, while HTR2A has been reported to be broadly expressed across germinal zones at later stages of human cortical development^20,21^, in our system its expression was more spatially enriched, suggesting temporally regulated serotonergic responsiveness. These differences likely reflect the earlier developmental stage modeled in assembloids and highlight the importance of considering developmental timing when interpreting neuromodulatory effects.

In addition, serotonergic innervation was not only associated with changes in basal progenitor populations but was also observed along the apical surface, where apical radial glia reside, suggesting that these cells may also be responsive to early neuromodulatory input. This effect appears to be selective, as intermediate progenitors did not show a comparable proliferative response, indicating subtype-specific sensitivity within cortical progenitor populations. Given the central role of apical progenitors in regulating early cortical expansion and lineage progression^46^, as well as evidence for species-specific regulation of their proliferative dynamics^17^, understanding how neuromodulatory signals influence these populations across developmental stages will be important.

Several limitations should be considered. While organoid-based systems capture key aspects of early human brain development^44^, they do not fully recapitulate the cellular diversity and environmental complexity of the in vivo brain^47^. In addition, although our data link serotonergic projections to changes in progenitor dynamics, the contributions of specific signaling pathways and interacting cell types remain to be resolved. At the same time, the modular and controllable nature of the assembloid system provides a key advantage, enabling targeted integration of region-specific inputs and perturbations to dissect neuromodulatory interactions in a defined human context.

Together, this work establishes a human assembloid platform to investigate early neuromodulatory input and positions serotonergic signaling as a modulatory influence on early cortical development in a controlled human model. This system provides a framework to define how serotonergic signaling shapes developmental trajectories and contributes to the emergence of sensitive developmental windows during which genetic or environmental perturbations may have lasting effects on the developing human brain.

## Acknowledgement

We thank Gina Tillmann and Helene Schamber for the pivotal technical support. We thank the DKFZ Single-Cell Open Lab (scOpenLab) for assistance with the scRNA sequencing experiments. We acknowledge the support of the NGS Core Facility Mannheim, Medical Faculty Mannheim of Heidelberg University.

## Declarations

### Funding

This work was generously supported by the Hector Stiftung II (to J.L.) and by the German Federal Ministry of Education and Research (BMBF) through the e:Med consortium SysMedSUDs (01ZX01909 to J.L., P.K.) and by the German Research Foundation (MA 4692/6-3 and MA 4692/12-1 to O.A.M)

## Competing Interests

The authors have no relevant financial or non-financial interests to disclose.

## Data and code availability

Raw data is available through the European Genome-Phenome Archive xxxx. All original code has been deposited at GitHub repository and is publicly available as of the date of publication (https://github.com/xxxx/xxxx).

## Author Contributions

Methodology: R.P.F., M.T.S., M.G., M.K., O.A.M.; Validation: A.A., L.W., C.G., A.H.; Formal analysis: R.P.F., M.T.S., M.G., M.K.; Investigation: R.P.F., M.T.S., M.G., M.K., A.A., L.W., C.G.; ScRNA-Seq data analysis, R.P.F., M.G., J.L.; Writing – Original Draft: R.P.F., M.T.S., M.G., A.A., J.L., Writing—reviewing and editing: all authors; Visualization: R.P.F., M.T.S., M.G., L.W.; Conceptualization: P.K., O.A.M, J.L.; Supervision: O.A.M., J.L; Funding acquisition: P.K., O.A.M., J.L.; Project administration: J.L.

---

**Supplementary Figure 1.**
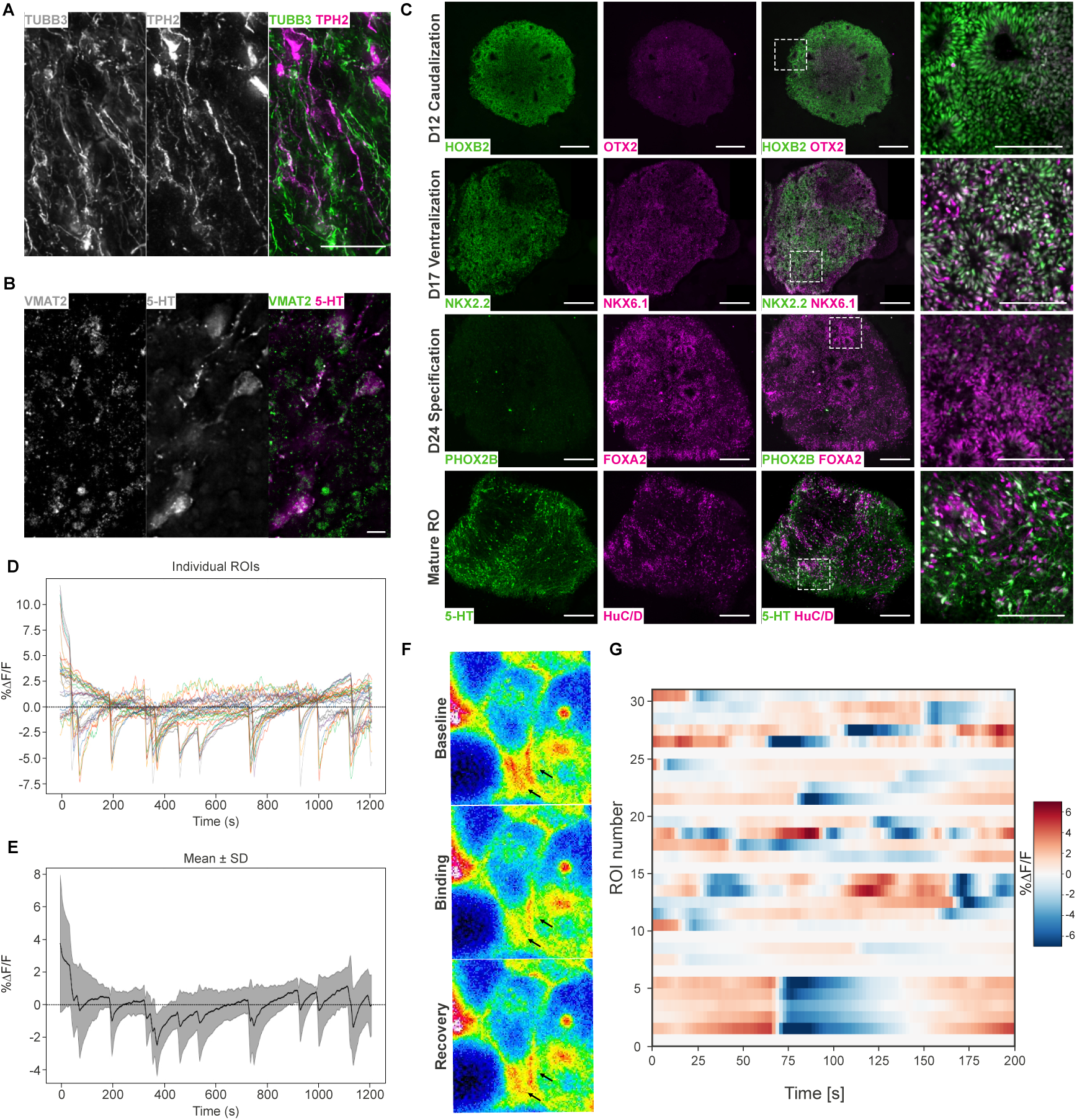
Characterization and functional validation of raphe organoids. **A** Representative immunofluorescence images showing expression of the neuronal marker TUBB3 (green) and the serotonin-synthesizing enzyme TPH2 (magenta), consistent with serotonergic neuronal identity. **B** Representative immunofluorescence images of raphe organoids (ROs) showing expression of the vesicular monoamine transporter VMAT2 (green) and serotonin (5-HT, magenta). **C** Representative immunofluorescence images of ROs from an independent iPSC line at sequential stages of differentiation, showing marker expression during caudalization (day 12; HOXB2, green; OTX2, magenta), ventralization (day 17; NKX2.2, green; NKX6.1, magenta), serotonergic lineage specification (day 24; PHOX2B, green; FOXA2, magenta), and neuronal maturation (day 60; 5-HT, green; HuC/D, magenta). White boxes indicate higher-magnification regions. **D** Quantification of sDarken fluorescence dynamics in ROs, shown as % ΔF/F traces of individual regions of interest (ROIs) over time. A total of 35 ROIs from 4 independent differentiations derived from 1 line were analyzed. **E** Mean % ΔF/F signal across ROIs (mean ± SD). **F, G** Characterization of sDarken responses in dissociated RO cultures. Representative images of sDarken-expressing cells during baseline, endogenous serotonin signaling, and recovery. Black arrows indicate regions exhibiting dynamic changes in fluorescence intensity (F). Heatmap of fluorescence changes (% ΔF/F) across 31 regions of interest (ROIs) over time from 6 independent batches derived from 1 line, showing dynamic fluorescence fluctuations consistent with endogenous serotonergic signaling (G). Scale bars: A, 20 µm; B, 50 µm; C, overview 200 µm; high magnification 100 µm.

**Supplementary Figure 2.**
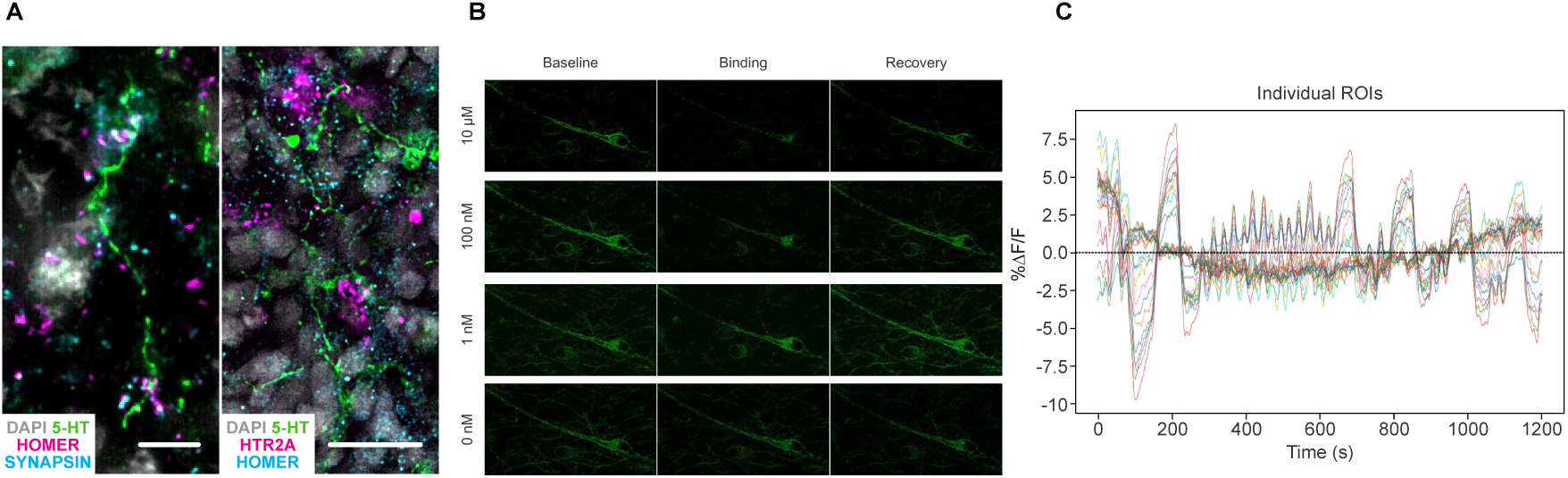
Synaptic features of serotonergic projections and functional validation of sDarken in cortical neurons. **A** Representative high-resolution immunofluorescence images showing serotonergic axons (5-HT, green) in cortical regions with punctate localization of synaptic markers SYNAPSIN (blue) and HOMER (magenta) at axon terminals (left), and showing expression of the serotonergic receptor HTR2A (magenta) and HOMER (blue; right). **B** Functional validation of sDarken in human cortical neurons. Representative images of sDarken-expressing neurons at baseline, during different concentrations of serotonin application (0 nM; 1 nM; 100 nM; 10 µM) and after washout and signal recovery. **C** Quantification of sDarken fluorescence dynamics in cortical regions of assembloids. A total of 22 regions of interest (ROIs) from 2 independent differentiations derived from 1 line were analyzed. Fluorescence changes are displayed as % ΔF/F traces of individual ROIs over time. Scale bars: A, 5 µm (left), 20 µm (right).

**Supplementary Figure 3.**
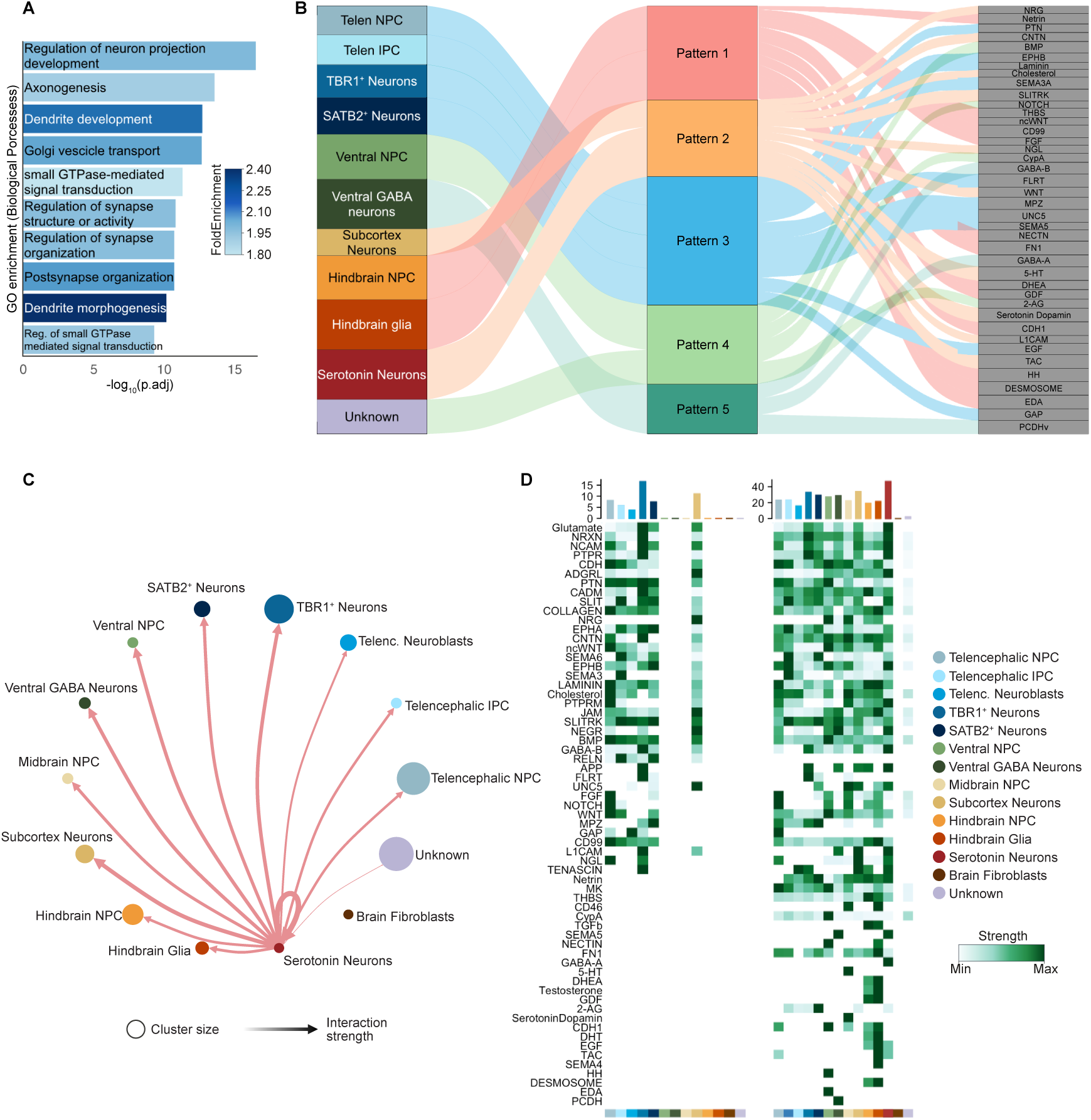
Transcriptional and intercellular signaling analyses in raphe–cortical assembloids. **A** Gene Ontology Biological Process (GO:BP) enrichment analysis of differentially expressed genes in TBR1⁺ neurons upon fusion. Top 15 biological processes are shown, ranked by adjusted p-value. **B** Pattern analysis of outgoing signaling networks inferred by CellChat. Signaling pathways are grouped into distinct patterns based on shared activity profiles across sender cell populations. **C** Network visualization of outgoing signaling from serotonergic neurons to other cell populations. Node size indicates the relative contribution of each cell population to the signaling network, and edge thickness represents inferred interaction strength. **D** Heatmap of incoming signaling patterns inferred by CellChat. Rows represent signaling pathways and columns represent receiver cell populations. Color intensity indicates relative incoming signaling strength. All snRNA-seq analyses shown in A–D are based on day 61 samples (9 organoids/assembloids; total n = 17,805 nuclei after quality filtering).

## Material and methods

### Culture of human iPSCs

Human induced pluripotent stem cell (iPSC) lines (B7_028#1, B7_028#4 from a 44-year-old female donor, 112#2 from a 19-year-old female donor and 35#2 from a 23-year-old male donor) derived from skin fibroblasts were used. All cell lines were obtained with written informed consent with approval of the Ethics Committee II of the Medical Faculty Mannheim of Heidelberg University (approval no. 2014-626N-MA). No analyses were performed to assess sex-specific effects.

iPSCs were maintained under feeder-free conditions on Geltrex (GT)-coated culture plates (Thermo Fisher Scientific) in Essential 8 (E8) medium at 37 °C, 5% CO₂, and ambient oxygen levels, with daily medium changes. Cells were passaged using EDTA (Thermo Fisher Scientific) and cultured at a ratio of approximately 1:3 to 1:10. After passaging, medium was supplemented with 5 μM Y-27632 (Cell Guidance Systems) for 24 h to enhance cell survival. All iPSC lines were routinely tested and confirmed negative for Mycoplasma spp. and were used at passages below 60. Pluripotency and genomic integrity of all lines were previously validated.

### Generation of raphe organoids

Human raphe organoids (ROs) were generated by stepwise patterning toward a caudal ventral hindbrain identity, followed by serotonergic specification. Human iPSC colonies were dissociated using TrypLE Express (Thermo Fisher Scientific) and seeded at 6,000-12,000 cells per well in 150 μL E8 medium supplemented with 50 μM Y-27632 into U-bottom 96-well plates pre-coated with 5% Pluronic F-127 (Sigma-Aldrich) in phosphate-buffered saline (PBS). At day 2, following aggregation, medium was switched to N2-based medium (DMEM/F12 supplemented with 1% B27, 0.5% N2 supplement, 1% GlutaMAX, 1% non-essential amino acids, 0.8 mg/mL D-glucose, 10 U/mL penicillin, 10 μg/mL streptomycin, and 50 μM 2-mercaptoethanol). For caudalization (day 2–12), medium was supplemented with 50 ng/mL FGF2, 7µg /mL insulin, 500 nM A83-01, and 200 nM LDN-193189, with medium changes every other day. At day 12, organoids were transferred to Pluronic F-127–coated dishes and cultured in ventralization medium consisting of N2-based medium supplemented with 50 ng/mL FGF2, 7 μg/mL insulin, 500 nM A83-01, 200 nM LDN-193189, 125 nM purmorphamine, and 250 nM smoothened agonist (SAG). Organoids were cultured on an orbital shaker at 70 rpm, 37 °C and 5% CO₂, with medium changes twice a week. At day 17, medium was replaced with specification medium (N2-based medium supplemented with 500 nM A83-01, 200 nM LDN-193189, 125 nM purmorphamine, 250 nM SAG, and 10 ng/mL FGF4. For neuronal maturation, organoids were cultured in Neurobasal medium supplemented with 0.5% N2 supplement, 1% B27, 1% GlutaMAX, 10 U/mL penicillin, 10 μg/mL streptomycin, 50 μM 2-mercaptoethanol, 0.8 mg/mL D-glucose, 2.5 μg/mL insulin, 200 μM L-ascorbic acid, 100 μM L-tryptophan, 10 ng/mL GDNF, 1 µM LM22A, 1µM LM22B, and 1 ng/mL TGF-β3. This medium was used as the standard maturation condition unless otherwise specified. During the first 14 days of maturation, 2.5 µM DAPT was added to promote neuronal differentiation. Medium was subsequently maintained without DAPT and changed twice weekly. After 14 days of maturation, organoids were transferred to a late maturation medium composed of Advanced MEM supplemented with 0.5% v/v N2 supplement, 2% v/v B27, 1% v/v GlutaMAX, 10 U/mL penicillin, 10 μg/mL streptomycin, 50 µM 2-mercaptoethanol, 800 µg/mL D-glucose, 2.5 µg/mL insulin, 200 µM L-ascorbic acid, 100 µM L-tryptophan, 10 ng/mL GDNF, 1 µM LM22A, and 1 µM LM22B.

### Generation of forebrain organoids

Forebrain organoids were generated following an established protocol^34–36^ with minor modifications. Human iPSC colonies were dissociated using TrypLE Express (Thermo Fisher Scientific) and seeded into low-attachment U-bottom 96-well plates pre-coated with 5% Pluronic F-127 (Sigma-Aldrich) in phosphate-buffered saline (PBS). Cells were plated at 6,000-12,000 cells per well in 150 μL E8 medium supplemented with 50 μM Y-27632, 10 U/mL penicillin and 10 μg/mL streptomycin. At day 5, neural induction was initiated by dual SMAD and WNT pathway inhibition using N2-based medium supplemented with 200 nM LDN-193189, 500 nM A83-01, and 2 µM XAV939, with medium changes every other day. Between days 9 and 11, upon formation of translucent neuroectodermal structures, organoids were embedded in GT (Thermo Fisher Scientific), transferred to differentiation medium as previously described^34–36^, and maintained under continuous orbital agitation at 70 rpm. Organoids were cultured at 37°C and 5% CO₂, with medium changes every 3–4 days. Between days 35 and 40, organoids were re-embedded in 2% w/v low-melting agarose (Sigma-Aldrich) and sectioned into 200-400 μm slices using a vibratome to enable long-term culture^35^. For sectioning, organoids were embedded in 4% w/v low-melting agarose (Carl Roth), and slicing was performed in ice-cold slicing medium. Slices were collected and maintained overnight in culture medium prior to further processing. Subsequently, slices were maintained under orbital agitation (70 rpm) in maturation medium consisting of differentiation medium supplemented with 1% v/v B27, 1µM LM22A, 1µM LM22B, 10 ng/mL GDNF, 200 µM L-ascorbic acid, and 0.2% v/v GT until further experimental use.

### Generation of raphe–cortical assembloids

Raphe (RO) and cortical (CO) organoids were generated separately according to their respective protocols and used for assembloid formation between days 35 and 45 of differentiation. Both organoid types were sectioned using a vibratome prior to assembly as descried above. For assembloid formation, individual RO and CO slices (200 μm) were transferred onto cell culture inserts (Merck) and positioned in close proximity to allow direct contact between tissues. Slices were maintained at an air–liquid interface^33^ under standard maturation conditions and cultured for at least three weeks, with daily medium changes. During this period, slices progressively adhered and formed stable assembloid structures.

### sDarken-based live imaging of serotonergic activity and analysis

Serotonergic activity was monitored using the genetically encoded serotonin sensor sDarken, as previously described^32^. sDarken (pLVX-EF1α-sDarken) was introduced via lentiviral transduction. For dissociated cultures, raphe organoids were enzymatically dissociated into single cells at day 24, transduced with lentivirus, and plated onto ECM-coated dishes in maturation medium supplemented with 5 µM Y-27632. Live imaging was performed 14 days after dissociation. For slice-based RO experiments, organoids were sectioned and transduced with sDarken at day 24. Slices were maintained in air–liquid interface culture as described above in late maturation medium and imaged after at least 3 weeks. For assembloid experiments, CO slices were transduced with sDarken 48 hours prior to fusion with raphe organoids, and assembloids were maintained in late maturation medium for at least 3 weeks. For validation of sensor functionality in a cortical context, human cortical neurons were generated from iPSCs via an NPC stage. NPCs were induced over 16 days using dual SMAD and WNT pathway inhibition–based conditions along established protocols^35^. Subsequently, NPCs were dissociated and differentiated into cortical neurons in defined media containing small molecules supporting neuronal maturation, as described previously^48,49^. Cells were cultured on Geltrex-coated dishes and maintained in differentiation conditions until use. Neurons were transduced with the serotonin sensor sDarken as described above. Fluorescence responses were assessed upon exogenous serotonin application (1 nM, 100 nM, and 10 μM). Prior to imaging, medium was replaced with imaging buffer containing 20 mM HEPES (pH 7.4), 140 mM NaCl, 2.5 mM KCl, 1.8 mM CaCl₂, 1 mM MgCl₂, and 10 mM D-glucose. Live imaging was performed using a Celldiscoverer 7 (Zeiss) at 37 °C and 4% CO₂. Images were acquired at 0.5 s intervals over a 5 min period. Regions of interest (ROIs) were drawn manually in ImageJ/Fiji around individual neurons, with one additional ROI designated as a background region. Mean grey values were extracted for each ROI across all frames and exported as CSV files. Background fluorescence was subtracted from each ROI trace by subtracting the background ROI value frame-by-frame. Fluorescence changes were expressed as ΔF/F, where baseline fluorescence (F₀) was defined as the median fluorescence of each ROI across the full recording duration. Traces were inspected visually and ROIs exhibiting movement artefacts or flat signals were discarded. Accepted traces were linearly detrended to remove slow <u>baseline</u> drift and subsequently smoothed with a Gaussian kernel (window = 5 frames, σ = 2.5 frames). Values were converted to % ΔF/F. All analysis steps were implemented in Python using a custom python package built on pandas, NumPy, and SciPy.

### Pharmacological treatment of cortical organoids

To assess receptor-specific effects of serotonergic signaling on cortical progenitor proliferation, cortical organoids (COs) were subjected to pharmacological perturbations at day 60. HTR2A signaling was activated using the selective agonist 720 nM TCB2, (Tocris Bioscience) and inhibited using the antagonist 252 nM ketanserin (ApexBio Technology) in the presence of 10 µM serotonin (Sigma-Aldrich). Control conditions included vehicle-treated organoids and treatment with serotonin alone. All treatments were performed for five days under standard culture conditions, with daily medium changes.

### Immunofluorescence

Organoid and assembloid samples were collected, washed in PBS, and fixed in 4% w/v paraformaldehyde (PFA) for 10–15 min at room temperature. After three washes in PBS, samples were incubated overnight at 4 °C in 30% w/v sucrose in PBS. Organoids were embedded in 7.5% w/v gelatin and 10% w/v D-sucrose, frozen in dry ice–cooled 98% v/v ethanol, cryosectioned at 20 μm, and mounted onto glass slides as descried^35^. Sections were permeabilized and blocked for 1 h at room temperature in PBS supplemented with 10% fetal bovine serum and 0.1–0.3% Triton X-100. Primary antibodies were applied overnight at 4 °C, followed by three washes in PBS and incubation with species-appropriate secondary antibodies for 1 h at room temperature. A complete list of antibodies, including sources and dilutions, is provided in **Supplementary Table 12**.

Nuclei were counterstained with DAPI, and samples were mounted using Mowiol. Image acquisition was performed using Leica confocal microscopes (TCS SP5 II and Stellaris 5) and a Leica DM6 B fluorescence microscope. Images were processed using Leica Application Suite software and Fiji (ImageJ).

### Immunofluorescence quantification

Quantification of immunofluorescence data was performed on organoid sections using region-specific analysis approaches. Images were acquired under identical settings within each experiment and analyzed using Fiji/ImageJ^50^ or CellProfiler^51^. For general marker quantification, marker-positive cells were quantified within manually defined regions of interest (ROIs) and expressed as the percentage of marker-positive cells relative to the total number of cells or, where indicated, relative to the total number of neurons. Multiple ROIs were analyzed per organoid, and values were averaged to obtain representative measurements for each sample. For basal progenitor analysis, mitotic basal progenitors were identified as pH3⁺ SOX2⁺ cells within the subventricular zone (SVZ), defined as the region basal to the ventricular zone (VZ). The proportion of proliferating basal progenitors was calculated relative to the total number of SOX2⁺ cells within the SVZ. Intermediate progenitor proliferation was assessed by quantifying pH3⁺ TBR2⁺ cells within the SVZ and expressed relative to the total number of TBR2⁺ cells. For assembloid analyses, mitotic activity within cortical regions was quantified by counting pH3⁺ cells within VZ and SVZ structures. Regions were classified based on the presence or absence of serotonergic projections, as identified by 5-HT immunoreactivity. Mitotic events were quantified within these regions and compared between innervated and non-innervated areas.

### Quantitative PCR (qPCR)

For each sample, 3–4 organoids were pooled to generate one biological replicate. Total RNA was extracted using the RNeasy Plus Mini Kit (Qiagen) according to the manufacturer’s instructions. cDNA was synthesized using the SuperScript III First-Strand Synthesis SuperMix (Thermo Fisher Scientific). Gene expression analysis was performed using gene-specific primers on a QuantStudio 7 Real-Time PCR System (Thermo Fisher Scientific). Primer sequences are listed in **Supplementary Table 13**. Ct values were normalized to the housekeeping gene RNA18S1 to obtain ΔCt values. Fold changes were calculated using the 2^−ΔΔCt^ method, using the preceding developmental time point as the reference for each comparison. Technical triplicates were averaged within each biological replicate prior to downstream analysis to avoid pseudo-replication. Log₂-transformed fold changes were used for statistical analysis, as specified in the corresponding figure legends.

### Sample preparation and single-nucleus RNA sequencing (snRNA-seq)

On day 61, after 21 days of culture at the air–liquid interface, samples were collected for single-nucleus RNA sequencing. Organoids were carefully lifted from membranes and transferred into microcentrifuge tubes. CO and RO control samples consisted of pooled organoids (n = 3 per sample). Assembloids (n = 3 per sample) were dissected using a scalpel to separate distinct regional compartments prior to processing, allowing independent representation of each tissue. Following removal of excess medium, samples were snap-frozen in liquid nitrogen and stored at −150 °C until further use.

Nuclei were isolated from samples using the Chromium Nuclei Isolation Kit with RNase inhibitor (10x Genomics, Pleasanton, CA, USA) according to the manufacturer’s instructions. Briefly, samples were mechanically homogenized, lysed, and filtered to obtain a purified nuclei suspension, followed by sequential washing and centrifugation steps to remove cellular debris. Nuclei concentration and quality were assessed using a LUNA-FL Dual Fluorescence Cell Counter. Approximately 10,000 nuclei per sample were loaded onto Chromium Next GEM chips for library preparation. snRNA-seq libraries were generated using the Chromium Next GEM Single Cell 3’ Kit v3.1 (10x Genomics) according to the manufacturer’s protocol. Libraries were sequenced on an Illumina NovaSeq 6000 platform (paired-end, 100 bp) at the Genomics & Proteomics Core Facility of the German Cancer Research Center (DKFZ).

### snRNA-seq data processing and analysis

FASTQ files were processed using Cell Ranger (10x Genomics) to generate filtered gene–barcode matrices. Downstream preprocessing was performed in R using Seurat v5^52^, unless otherwise stated. Filtered matrices were imported using Read10X and converted into Seurat objects. Per-cell quality control metrics were computed, including the number of detected genes, total UMI counts, and the fraction of mitochondrial transcripts. Cell filtering was performed programmatically within the preprocessing workflow using a robust, sample-wise approach based on median absolute deviations (MADs). For each sample independently, nFeature_RNA and nCount_RNA were log10-transformed (log10(x + 1)), and cells were retained only if both metrics fell within the median ± 3 MADs of the corresponding distribution. In addition, cells with >5% mitochondrial transcripts were excluded. For cross-platform integration, filtered count matrices together with feature, barcode, and metadata tables were exported and reconstructed in Python using Scanpy^53^. Highly variable genes were recomputed using scanpy.pp.highly_variable_genes with the cell_ranger flavor, selecting the top 2,000 genes, with sample identity specified as a batch variable. Only highly variable genes were retained for integration. Latent space integration was performed using scVI^54^, with raw counts as input and sample identity specified as a categorical batch covariate. Models were trained using GPU acceleration with mini-batch optimization (batch size = 128) for up to 400 epochs. The learned latent representation was used to construct a nearest-neighbor graph (n_neighbors = 15) and to compute a UMAP embedding. Cell type annotation was performed using a combination of automated prediction and manual curation. Normalized expression data were analyzed using CellTypist^37^ with the “Developing_Human_Brain”^38^ reference model and majority voting enabled. Predicted labels and confidence scores were incorporated into the metadata and used as a first-pass annotation. Final cell identities were assigned based on concordance between CellTypist predictions and canonical marker gene expression, followed by manual refinement. Downstream analyses were performed in Seurat. Samples were grouped into three experimental conditions (COs, ROs, and RO-CO assembloids). Marker expression across annotated populations was visualized using DotPlot. Cell-type composition was quantified by calculating the proportion of cells per condition. Cell cycle position analysis was performed using tricycle. The Seurat object was subset to telencephalic neural progenitors from COs and RO-CO assembloids, and tricycle positions were computed on the RNA assay using human gene symbols as input features. Condition-specific distributions along the inferred cell cycle continuum were visualized using kernel density estimation. Differential expression analysis was performed using FindMarkers on a combined identity class (condition × cell type), comparing RO-CO assembloids versus COs within matched telencephalic populations. Functional enrichment analysis was performed on genes differentially expressed in RO-CO assembloid compared to their control counterparts. Significant genes were filtered (adjusted p-value < 0.05), ranked by effect size, and ribosomal genes were excluded prior to enrichment analysis. Gene symbols were converted to Entrez identifiers and subjected to Gene Ontology enrichment analysis using clusterProfiler^55^ for Biological Process terms with Benjamini–Hochberg correction. Enriched terms were visualized using bar plots scaled by fold enrichment. Gene–term associations were extracted from enrichment results and combined with differential expression statistics to generate gene-category networks for downstream visualization.

Cell–cell communication analysis was performed using CellChat^40^. Normalized expression data were grouped by annotated cell type, and a CellChat object was constructed using the human ligand–receptor interaction database. Overexpressed genes and interactions were identified, communication probabilities were computed, and ligand–receptor interactions were aggregated into pathway-level communication networks. Outgoing and incoming signaling strengths were quantified using network centrality analysis and used to compare communication patterns across conditions. All analyses were performed using Seurat, Scanpy, scvi-tools, CellTypist, clusterProfiler, and associated R and Python packages. Scripts and computational environments are described in the Data and Code Availability section.

## Statistical analysis

Quantification and statistical analyses were performed in R (v4.4.2) using custom scripts together with the readxl, rstatix, ggpubr, ggplot2, and scales packages. For qPCR analyses, ΔΔCt values were calculated using the preceding developmental time point as reference, and fold changes were computed as 2^-ΔΔCt^. Statistical testing was performed on log2-transformed fold changes using pairwise t-tests between adjacent differentiation stages (D0 versus D12, D12 versus D17, and D17 versus D24), followed by Benjamini–Hochberg correction. For rostro-caudal patterning and serotonergic differentiation analyses, marker-positive fractions were calculated as percentages relative to DAPI-positive nuclei and HuC/D-positive neurons, respectively. For proliferation analyses in fusion experiments, comparisons between innervated and non-innervated VZ/SVZ regions were performed using Wilcoxon rank-sum tests with Benjamini–Hochberg correction. For pharmacological perturbation analyses in the SVZ-like bRG compartment, the proliferative fraction was calculated as pH3⁺/SOX2⁺ × 100. Treatment labels were standardized, and measurements were averaged within each combination of condition, batch, and sample prior to testing. Pairwise comparisons across treatments were performed using Wilcoxon rank-sum tests with Benjamini–Hochberg correction. For intermediate progenitor analyses, pairwise comparisons between conditions were performed using Wilcoxon rank-sum tests with Benjamini–Hochberg correction. For all assays, the experimental unit was defined according to the aggregation step implemented for each dataset, and measurements derived from the same biological unit were averaged prior to statistical testing to avoid pseudoreplication. Statistical inference was therefore performed on aggregated values. Data were visualized as boxplots overlaid with jittered individual points, with point shape indicating cell line. In all boxplots, the center line represents the median, the box indicates the interquartile range (25th–75th percentiles), and whiskers extend to 1.5 × the interquartile range. Individual points correspond to aggregated observations after the assay-specific summarization step. All statistical tests were two-sided unless otherwise specified, and P values were adjusted using the Benjamini–Hochberg procedure where applicable. Adjusted P values were displayed directly on the plots, and non-significant comparisons were omitted from annotation.

## Notes

### Competing Interest Statement

The authors have declared no competing interest.

